# Spider lures exploit insect preferences for floral colour and symmetry

**DOI:** 10.1101/693648

**Authors:** Thomas E. White, Darrell J. Kemp

**Affiliations:** School of Life and Environmental Sciences, The University of Sydney, Sydney, Australia 2106; Department of Biological Sciences, Macquarie University, North Ryde, Australia 2113

**Keywords:** sensory trap, sensory bias, orb-web spider, prey lure, mimicry

## Abstract

Sensory systems can capture only a fraction of available information, which creates opportunities for deceptive signalling. The sensory traps and sensory bias models have proven valuable for explaining how visual systems and environments shape the design of sexual signals, but their application to deceptive signals is largely untapped. Here we use the ‘jewelled’ orb-web spider *Gasteracantha fornicata* to experimentally test two longstanding hypotheses for the function of deceptive visual lures. Namely, that they: (1) exploit generalised preferences for conspicuous colouration (sensory bias), or (2) co-opt the otherwise-adaptive foraging response of prey toward flowers (sensory traps). In a field-based study we manipulated the conspicuous dorsal signal of female spiders along two axes —– colour pattern and symmetry — to generate a gradient of floral resemblance, and monitored the per-individual consequences for prey interception. As predicted by the traps model, the most attractive phenotypes were those with flower-like radial symmetry and solid colour patterns, and their attractiveness equaled that of wild-type models. These results demonstrate that deceptive orb-web spider lures function, in part, as inter-kingdom sensory traps via floral mimicry, and support the broader extension of sensory-based models to deceptive signalling contexts.

## Introduction

Visual communication is ubiquitous, and the demands of effective information-exchange have driven diverse outcomes (Maia et al. 2013; Thoen et al. 2014; Dalrymple et al. 2015). Understanding this diversity requires examining the relationship between signals, environments, and sensory systems, for which the sensory traps and bias models — under the umbrella of sensory drive — have proven valuable (among a suite of related models; Christy 1995; Endler 1992; Endler and Basolo 1998; West-Eberhard 1979). According to the sensory trap model, signals evolve under a model-mimic dynamic to co-opt receiver responses that function adaptively in otherwise unrelated behavioural contexts (Christy et al. 2003). This model accounts for how the design of sexual signals, for example, may be shaped by how potential mates detect or recognize food items (Rodd et al. 2002) or shelter (Christy et al. 2003). The sensory bias model, by contrast, emphasizes how underlying sensory and/or perceptual biases may present opportunities for exploitation and hence drive signal evolution (Basolo and Endler 1995; Ryan and Cummings 2013). The elaborate fins of male swordtails present a canonical example (Basolo 1990), having evolved in response to a pre-existing female bias toward such structures (Basolo 1990; Basolo 1995). Each of these two models has robust empirical support in the context of sexual signalling, however much remains to be learned about their ability to explain signal evolution more broadly.

Visual luring is a widespread predatory strategy and is particularly common among sit- and-wait predators. Orb-web spiders are a model group, with many species combining striking body colours and patterns to actively attract insect prey to the web (Tso et al. 2004; Chuang et al. 2007a; White and Kemp 2015). The question of why such conspicuous deceptive signals are attractive to insect viewers has been the focus of considerable attention (Tso et al. 2004; Chuang et al. 2007b; Rao et al. 2015; Goncalves and Gawryszewski 2017; White and Kemp 2017). Two hypotheses predominate, which informally mirror the bias and traps models; namely, that lures (1) exploit innate colour preferences, or (2) co-opt the foraging response of prey toward flowers. Empirical support for these hypotheses is presently limited to observational and correlative data, and hence remains equivocal (e.g., Tso et al. 2004; Chuang et al. 2007b; Goncalves and Gawryszewski 2018; White et al. 2017). Formalising these hypotheses within the models of sensory theory offers a promising path to progress and may prove reciprocally beneficial in guiding future studies of deceptive signalling.

Whereas predictions from the bias and traps models overlap to some degree, their core predictions as applied to deceptive lures can be neatly partitioned (White & Kemp 2015). If conspicuous visual lures are exploiting receivers’ sensory biases, then the most likely perceptual target is colour. The insect prey of luring predators are taxonomically diverse, albeit with an overrepresentation of pollinating flies and bees (Nentwig 1985; Nentwig 1987; O’ Hanlon et al. 2014a). Strong innate preferences for (human-perceived) yellows and whites are well documented (Kay 1976; Lunau 1988; Lunau and Maier 1995), which parallels a notably biased distribution of these colours among predator lures (White and Kemp 2015). A standing prediction under the bias model, then, is that the expression of preferred colours among deceptive signallers should predict their attractiveness to potential prey. The traps hypothesis, by contrast, suggests that lures are exploiting an otherwise-adaptive attraction to flowers in a dynamic more closely akin to floral mimicry. When foraging, pollinating insects integrate the aforementioned colour preferences with information on different forms of floral symmetry, which they can readily perceive and express preferences for (as contrasted with asymmetry, which is aversive: Chittka and Raine 2006; Kay 1976; Lehrer et al. 1995; Lunau and Maier 1995; Giurfa et al. 1996), and radial symmetry is both the most ancient and common form showcased among angiosperms (Crane et al. 1995; Neal et al. 1998; Endress 2001). Accumulating evidence for the predicted resemblance between lures and these features of sympatric flowers supports this mimetic view (Tso et al. 2004; Goncalves and Gawryszewski 2017; White et al. 2017). The key untested prediction, however, is that lures should co-opt the prey’ s natural response toward flowers. The strongest evidence to date comes from the orchid mantis, which resembles sympatric flowers and presents a more attractive signal to pollinators (O’ Hanlon et al. 2013; O’ Hanlon et al. 2014a; O’ Hanlon et al. 2014b). Although this presents a compelling example of pollinator deception, the restricted range of experimental stimuli offered to viewers in the key assay presents a challenge to unambiguously distinguishing between the traps and (more permissive) bias explanations.

Here we sought to formalise and test these adaptive hypotheses for deceptive signalling using the jewelled orb-web spider *Gasteracantha fornicata* (supplementary Fig. S1). Females of the species are colour polymorphic sit-and-wait predators, whose striking yellow- or white-and-black banded abdomens lure prey — primarily pollinating Diptera and Hymenoptera — to their webs (Hauber 2002; Kemp et al. 2013; White and Kemp 2016). To distinguish between the traps and bias hypotheses we manipulated the appearance of wild female *G. fornicata* in their natural habitats along two independent axes — colour and symmetry (Fig. 1). Our manipulations consisted of nine different treatments (including the wild-type) that encompassed the full-factorial combination of three levels of colour and three levels of symmetry. The sum of treatments represented an approximate gradient of floral resemblance, thereby affording clear predictions for relative attractiveness under a (generalized) sensory trap hypothesis (i.e., the x-axis of Fig. 2). Predicted attractiveness under the sensory bias model is however different because the main vector of attractiveness in this case should relate to stimulus color alone (the y-axis of Fig. 2). We evaluated these predictions according to realized prey capture rates of wild, free-ranging spiders randomly assigned among the nine treatment stimuli.

**Figure 1:**
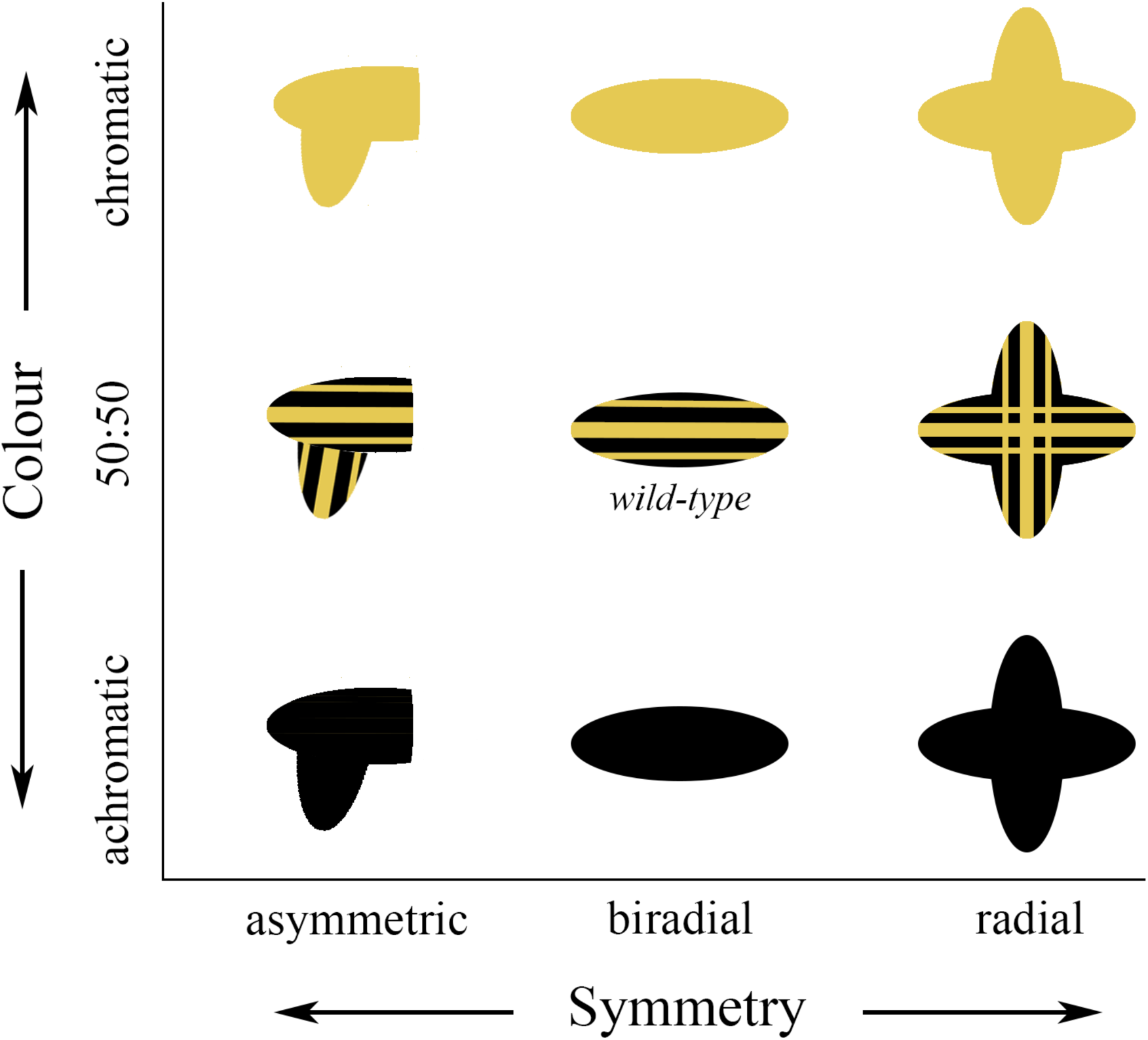
The colour pattern manipulations as applied to naturally-occurring female specimens of *G. fornicata*. The aim was to represent an approximate gradient of floral resemblance from most flower-like (top right) to least (bottom left), while including a wild-type model (center; also see supplementary fig. S1).

**Figure 2:**
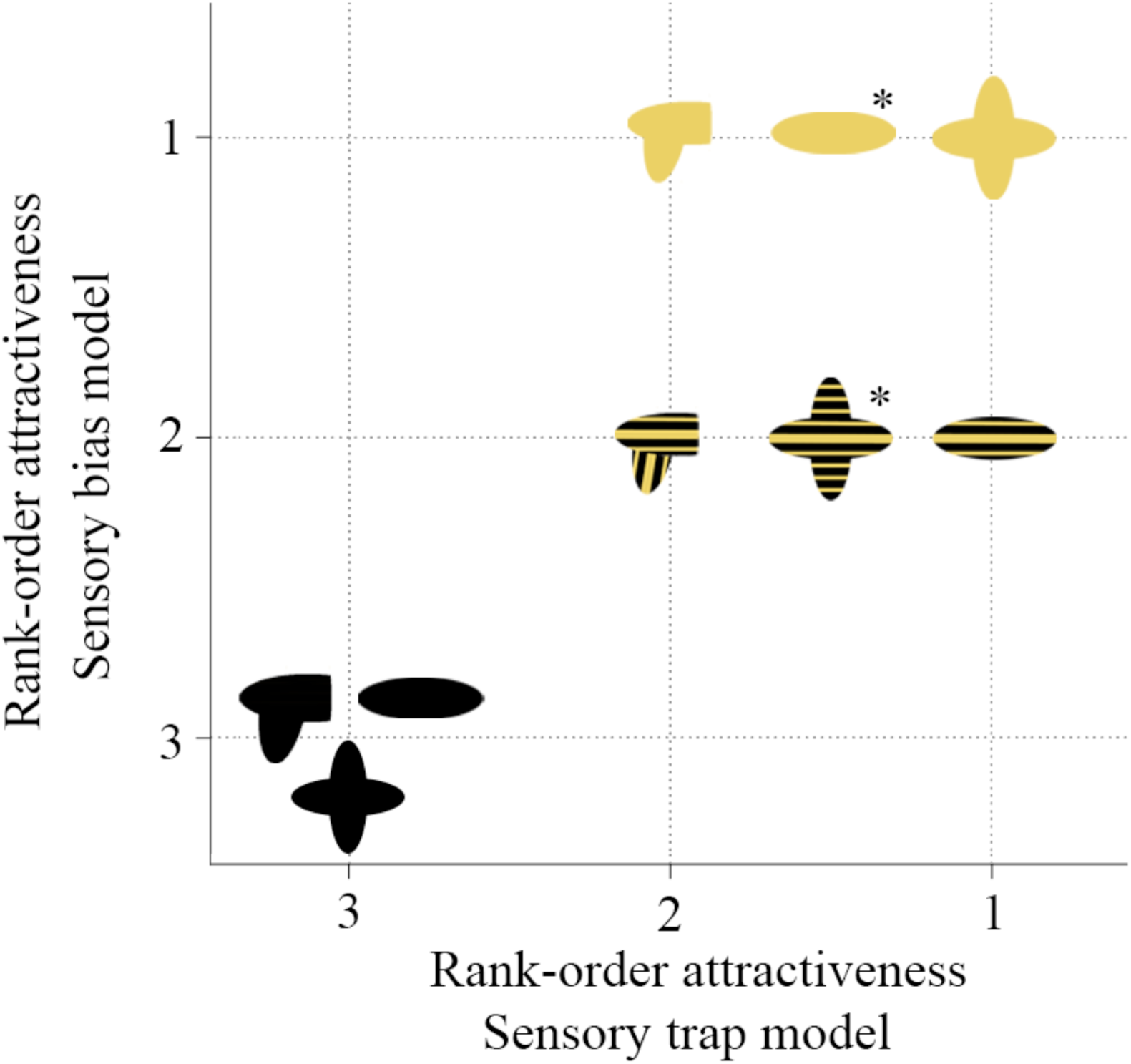
The predicted attractiveness of artificial phenotypes under the traps and bias models of signal evolution. If lures are exploiting general colour-biases, the attractiveness of models should simply be predicted by yellow coverage. If, in contrast, lures are floral sensory traps, then the natural phenotype should be as equally attractive as the most ‘flower-like’ phenotype. Note that solid biradial and striped radial models (asterisked) are of intermediate rank on the x axis because it is difficult to derive unambiguous predictions for their rank-order attractiveness under the sensory traps model.

## Methods

### Phenotype manipulations and prey interception rates

Our manipulative treatments included asymmetric, biradially symmetric, and radially symmetric shapes, in a fully factorial combination of solid black, black-and-yellow banded, and solid yellow patterns (n = 17-29 each; Fig. 1). We manipulated the appearance of spiders by fixing a painted cardboard model (Quill 180 gsm paper) corresponding to a given treatment (Fig. 1) to each individual’s otherwise flat dorsal abdomen using a ca. 5 mm^2^ square of double-sided tape. Importantly, we controlled the proportionate size of stimuli in each symmetry-class to ensure an equal area of colour coverage. That is, all solid-yellow models displayed approximately the same total amount of yellow (ca. 81 mm^2^), all striped models had equal amounts of yellow and black (ca. 40 mm^2^), and all black models displayed the same amount of black (ca. 81 mm^2^). We used Derivan Matisse Yellow-Mid AZO Series 2 paint to imitate the yellow colouration of *G. fornicata*, which has previously been spectrally matched for this purpose using standard methods and is also a known match to sympatric insect-pollinated flora (Maia et al. 2019; White and Kemp 2017). In addition to the nine primary treatments we included a further control in which spiders were unmanipulated save for a square of double-sided tape on their ventrum. Although *G. fornicata* are colour polymorphic, we used only yellow colouration in all treatments for simplicity and manipulated both ‘white’ and ‘yellow’ individuals in the field. There is some evidence for microhabitat differentiation between *G. fornicata* morphs (White and Kemp 2016), but our application of treatments was randomised and hence simply contributes unbiased residual variation (i.e., noise). The extent of any microhabitat effects therefore adds conservatism to our focal contrasts.

To estimate prey interceptions as a key component of fitness we used a transect-based method comparable to one previously used in this system (White 2017). After applying the cardboard models, we recorded the presence of new prey and/or web damage at 30 minute intervals for 4 hours, either in a morning (0800-1200) or, less often, afternoon (1300-1700) session. Abiotic confounds (such as web damage by wind-blown debris) may inflate true interception rates, but such effects would again be randomly distributed across treatments and simply inflate residual variation. Spiders whose webs that sustained > 50% damage during an observation period were taken to indicate gross environmental disturbance and were excluded (n = 12) as well as those whose model did not remain affixed (n = 4). All work took place in November 2018 across populations spanning Cairns to Port Douglas, Queensland, Australia. The observer (TEW) could not possibly be blind in regard to treatments, but the unambiguous response variable should work to ameliorate unconscious bias.

### Statistical analyses

To validate the baseline efficacy of the phenotypic manipulations, we first tested for differences in prey interceptions between the wild-type models of *G. fornicata* (biradial striped; Fig. 1 centre) and unmanipulated spiders using a generalised linear mixed-effects model (GLMM). We specified interception rate (mean interceptions / 30 minutes) as the Gaussian response following confirmation of the normality within groups, and treatment (presence/absence) as a main effect, with diel session (morning/afternoon) as a random covariate to account for any systematic differences associated with diel insect activity.

For the central tests we used a GLMM with interception rate (mean interceptions / 30 minutes) as the response, as above. We specified an interaction between colour (black/striped/solid) and symmetry (asymmetric/biradial/radial) and their main effects, and included diel session (morning/afternoon) as a random covariate. We then used Tukey post-hoc contrasts to test for pairwise differences across all treatment combinations. Should the sensory bias model best explain the attractiveness of phenotypes we predict a main effect of colour alone (Fig. 2a). In contrast, the sensory traps hypothesis predicts an interaction between colour and symmetry, with post-hoc tests revealing grouped differences in the manner specified in Figure 2b (and as discussed above). Summary statistics reported below are pooled means ± standard deviations of prey interceptions rates (interceptions / 30 minutes). All analyses were run in R v. 3.5.2 (R Core Team 2018) using ‘nlme’ (Pinheiro et al. 2018) for linear mixed modelling and ‘multcomp’ (Hothorn et al. 2008) for multiple comparisons.

### Data availability

All data and code will be made persistently available via Github and Zenodo upon acceptance.

## Results

We found no difference in prey interception rates between control *Gasteracana. fornicata* and wild-type models (F_1, 41_ = 0.65, *p* = 0.43, R^2^ = 0.02). The vanishingly small effect size between each group moreover supports the absence of any biologically-relevant consequence of handling. For the main test, we found an interactive effect of colour and symmetry on prey interception rates (F_4,218_ = 4.12, *p* = < 0.01, conditional R^2^ = 0.54), as well as main effects of colour (F_2,218_ = 107.40, *p* = < 0.01) and symmetry (F_2,218_ = 15.08, *p* = < 0.01). Pairwise contrasts (supplementary table S1) revealed considerable variation in prey interception rates between treatments, with three distinct phenotypic groupings (Fig 3). Spiders assigned to black control treatments intercepted prey less frequently than all others (0.84 ± 0.77), while both striped- and solid-coloured asymmetric phenotypes had greater capture success (1.92 ± 0.70). The highest rates of prey interception were shared by radially and biradially symmetric treatments across both striped- and solid-coloured phenotypes (2.86 ± 0.89)

**Figure 3:**
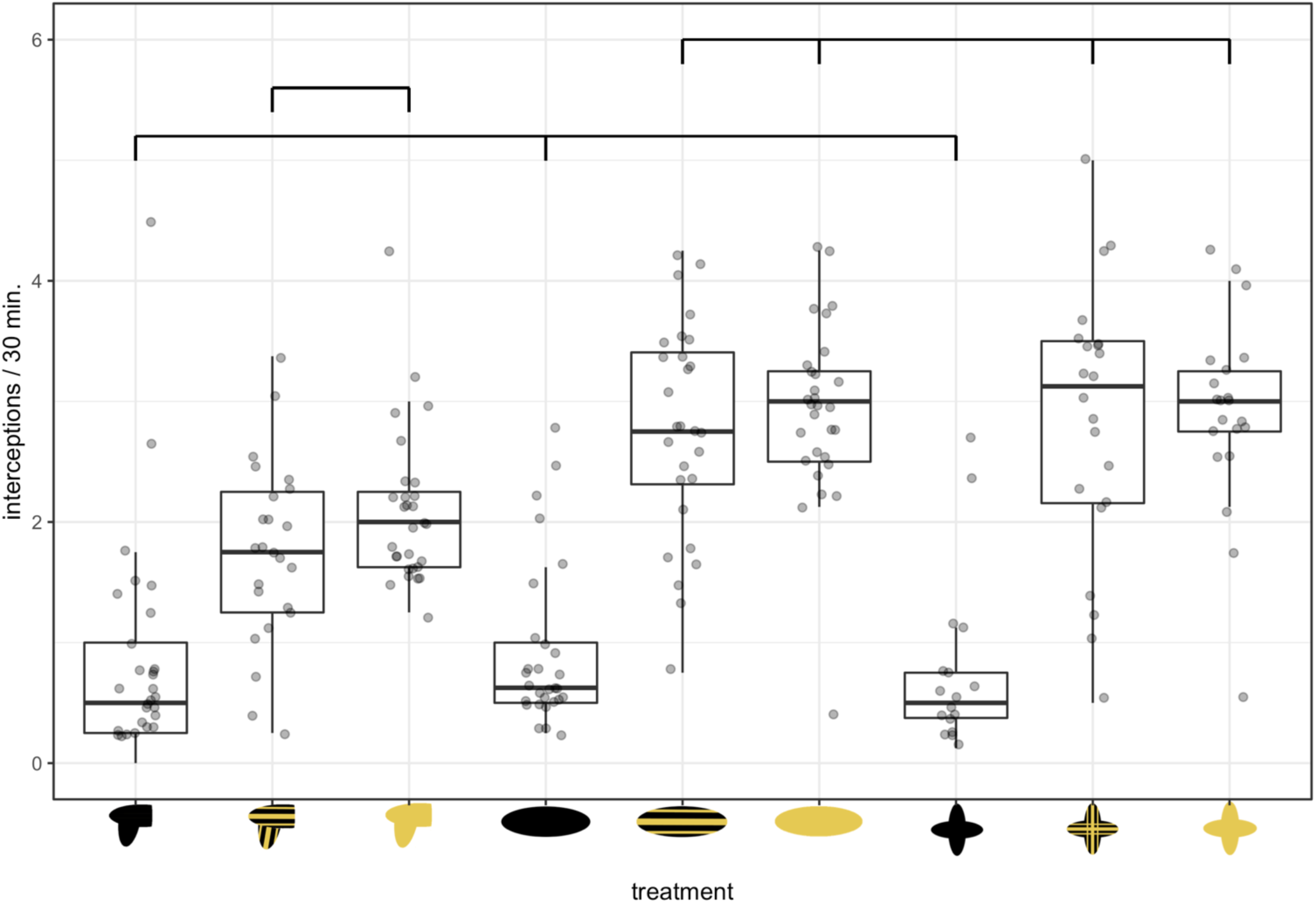
The effect of phenotypic manipulations (Fig. 1) on prey interception rates in *G. fornicata*. Boxes denote the median and first and third quartiles, while whiskers extend to values at a maximum of 1.5 times the inter-quartile range. Horizontal lines indicate statistically distinct treatment groupings based on post-hoc multiple comparisons. Sample sizes, left-to-right; 29, 24, 29, 29, 28, 29, 17, 22, 21.

## Discussion

Visual lures are a striking adaptation for predation, but the mechanism of deception is poorly resolved. Here we manipulated the phenotypes of the jewelled spider *Gasteracantha fornicata* along an approximate gradient of floral resemblance to test whether deceptive lures are exploiting simple colour-biases, or co-opting foraging preferences, in prey. As predicted by the sensory traps model (Fig. 2, x axis), we found equivalently heightened prey interception rates between the natural phenotype and the biradially symmetric, solid-yellow (most ‘floral’) model (Fig. 3). In contrast, the sheer coverage of yellow colouration on models was not solely predictive of prey interceptions as expected under a bias explanation (that is, the effect of colour manifested via an interaction with symmetry). Control tests suggest that the manipulations were effective and highly specific in consequence, with no difference in interception rates between the wild-type model and unmanipulated spiders, and significant differences between black models and all others (Fig. 3; supplementary table S1). In sum, our results suggest that female *G. fornicata* co-opt the foraging responses of prey toward flowers, in a deceptive inter-kingdom sensory trap.

Though the wild-type and most ‘flower-like’ phenotypes were equally attractive (Fig. 3), *Gasteracantha fornicata* are unlikely to be a simple mimic of any one sympatric flower. Rather, the signals of spiders are likely presenting a combination of visual cues that are shared by local flowers including, but not limited to, the spectral, spatial, and symmetric properties of patterns (O’Hanlon et al. 2014a; O’Hanlon et al. 2014b; White et al. 2017). This accords with known features of visual processing among well-studied insects in which local cues such as (in order of prioritisation) colour, modulation, shape, area, and orientation are weighed and integrated to guide the choice and classification of stimuli (Giurfa et al. 1995; Horridge and Zhang 1995; Giurfa et al. 1996; Horridge 2007). These cues can be readily generalised to novel contexts (Stach et al. 2004), and their relative importance may vary during assessments of mimetic accuracy (e.g. colour similarity may prioritised over shape; Kazemi et al. 2014) or with cognitive load (e.g. in speed-accuracy tradeoffs; Chittka & Osorio 2007). This offers a basis for deceptive signal efficacy among luring predators despite their human-subjective distinctiveness from flowers. That is, lures may need only present an ensemble of a few salient cues, rather than a faithful analogue of floral signals, to exploit the foraging response of insect prey (discussed further below). This possibility is further enabled by the phenotypic diversity of sympatric flora, which present a suite of shapes, symmetries, and colour patterns from which deceptive signallers may draw (see White et al. 2017 for data relevant to *G. fornicata* specifically). Our finding that colour alone was attractive to insects, yet moreso when combined with floral symmetry cues, is consistent with such a view (Fig. 3), though awaits closely controlled behavioural work to test in detail.

While the presence of colour in any form was associated with improved attractiveness, the colour pattern —– be it solid or striped — had no further effect (Fig. 3). There are two plausible explanations for the lack of a pattern effect. One is that the stripes cannot be resolved at meaningful distances, and a striped pattern would instead only generate a subtly duller, though still ‘solid’ signal that is functionally equivalent to their block-coloured counterparts. Although the stripes are indeed likely to be resolved only at close distances by typical fly and bee viewers (Land 1997), past work has shown that interception rates are directly modified by the orientation of the stripes of *G. fornicata* in the web (White 2017), thereby establishing the discriminability of the patterns at relevant viewing distances. A simple alternative, related to the above, is that both striped and solid variants present attractive cues to viewers that are shared by flowers. Solid colours are typical among flowers, though some 33% of radially symmetric and 14% of bilaterally symmetric species also present patterned ‘floral guides’ (Dafni and Giurfa 1999). Such guides take the form of repeated stripes and/or radiating elements, which serve to draw pollinators to the location of nectar and pollen centers (Dafni and Kevan 1996; Dafni and Giurfa 1999). The banded pattern of *G. fornicata* and our striped, radial model are thus unlikely to be entirely novel to experienced receivers and may merely present another cue that pollinators recognise as broadly ‘floral’.

The role of colour in visual deception is widespread, and our results support the extension of sensory models to formalise the study of its causes and predicted consequences more generally. The dynamic displays of crab spiders (Heiling et al. 2003), red rims of pitcher plants (Schaefer and Ruxton 2008), and decorated webs of spiders (Herberstein et al. 2000) are striking examples, though identifying the underlying mechanism in each case has proven difficult (Herberstein et al. 2000; Schaefer and Ruxton 2009). Our results reiterate the well understood necessity of considering the perspective of receivers, since human-subjective assessments of similarity are a poor guide to the existence and extent of mimicry (Fig. 3). Though our wild-type and ‘floral’ spider models bear little human-subjective resemblance, our results are consistent with the view that they converge at some stage of sensory processing in insect viewers to elicit a shared foraging response (as noted above). This accords with evidence from sexual signalling systems in which the co-option of food detection pathways underlies the attractiveness and early evolution of male sexual ornaments, such as the yellow caudal bands of male swordtail characins (Garcia & Ramirez 2005; Rodd et al. 2002). Interestingly, once such signals become common within a population, receivers may ‘escape’ the sensory trap via selection for increased response thresholds or improved discriminability (Garcia & Ramirez 2005). We may predict a similar course in luring systems, though the consequences for signal evolution will diverge due to differences in the alignment of interests between signallers and receivers. In sexual contexts the interests of both parties are broadly aligned toward reproduction. Although selection may favour the partitioning of receivers’ feeding and sexual responses through improved discrimination of mimetic traps, they will ultimately respond positively to both sexual and foraging cues (Basolo and Endler 1995; Ryan and Cummings 2013). With respect to signallers, a known consequence is a shift toward signal honesty which also reduces the foraging costs to receivers of responding to deceptive cues (Garcia & Ramirez 2005). Luring systems, in contrast, cannot follow such a trajectory since they are entirely antagonistic. Thus while selection for improved discrimination and response thresholds in receivers is a predictable outcome, the consequences for deceptive, as opposed to sexual, signal evolution will diverge. Possible outcomes include selection for improved mimetic fidelity via the integration of new cues and/or refinement of existing ones (e.g. a move toward closer spectral or morphological resemblance to models), a shift toward dietary specialisation or generalisation depending on the composition of available prey (and their foraging preferences), and/or the evolution of signal polymorphism if available prey and models are diverse enough to generate multiple fitness optima (Kazemi et al. 2014; Kikuchi & Pfenning 2013; White & Kemp 2016). These are intriguing avenues for future work and highlight the reciprocal promise of luring systems for fueling both empirical insight and theoretical development.

## Acknowledgments

TEW thanks Elizabeth Mulvenna and Cormac White for their endless support.

