## Supplementary material for "Spider lures exploit insect preferences for floral colour and symmetry"

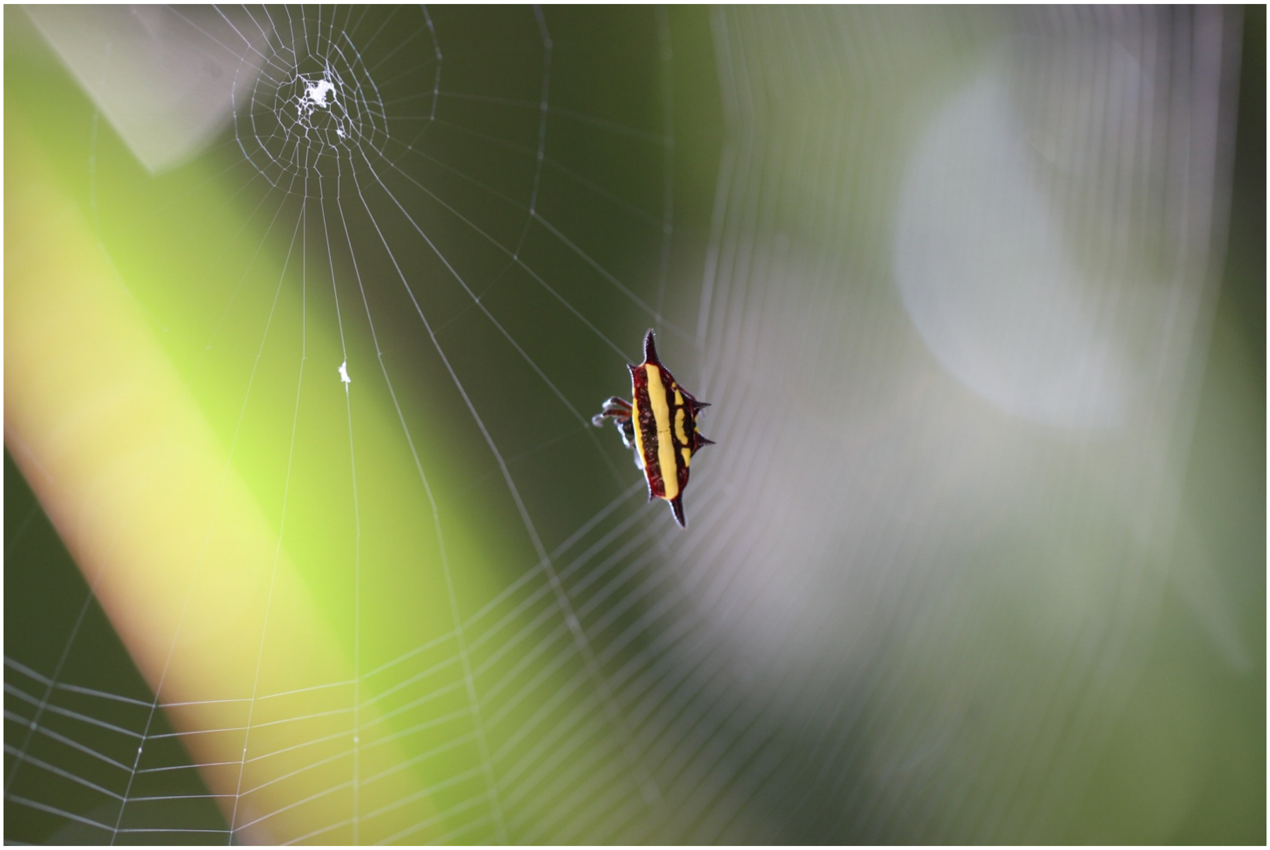

**Supplementary Figure S1:** Female *Gasteracantha fornicata* from Cairns, QLD, Australia, weaving a web (photograph: Thomas E. White).

**Supplementary Table S1:** The results of post-hoc contrasts testing the effects of colour and symmetry on prey interception rates in *G. fornicata.* Contrasts detail pairwise combinations of black, striped, and yellow models across radial (rad), biradial (birad) and asymetrical (asym) treatments.

| ***Contrast*** | ***Estimate*** | ***Std. Err.*** | ***z*** | ***P*** |
|---|---|---|---|---|
| stripe.asym - black.asym | 0.89 | 0.23 | 3.94 | 0.00 |
| yellow.asym - black.asym | 1.24 | 0.21 | 5.77 | < 0.001 |
| black.birad - black.asym | 0.06 | 0.21 | 0.26 | 1.00 |
| stripe.birad - black.asym | 1.91 | 0.22 | 8.84 | < 0.001 |
| yellow.birad - black.asym | 2.07 | 0.21 | 9.66 | < 0.001 |
| black.rad - black.asym | -0.08 | 0.25 | -0.33 | 1.00 |
| stripe.rad - black.asym | 2.00 | 0.23 | 8.67 | < 0.001 |
| yellow.rad - black.asym | 2.05 | 0.23 | 8.75 | < 0.001 |
| yellow.asym - stripe.asym | 0.35 | 0.23 | 1.56 | 0.83 |
| black.birad - stripe.asym | -0.83 | 0.23 | -3.69 | 0.01 |
| stripe.birad - stripe.asym | 1.02 | 0.23 | 4.51 | < 0.001 |
| yellow.birad - stripe.asym | 1.18 | 0.23 | 5.25 | < 0.001 |
| black.rad - stripe.asym | -0.97 | 0.26 | -3.74 | 0.01 |
| stripe.rad - stripe.asym | 1.11 | 0.24 | 4.62 | < 0.001 |
| yellow.rad - stripe.asym | 1.16 | 0.24 | 4.76 | < 0.001 |
| black.birad - yellow.asym | -1.18 | 0.21 | -5.51 | < 0.001 |
| stripe.birad - yellow.asym | 0.67 | 0.22 | 3.11 | 0.05 |
| yellow.birad - yellow.asym | 0.83 | 0.21 | 3.88 | 0.00 |
| black.rad - yellow.asym | -1.32 | 0.25 | -5.29 | < 0.001 |
| stripe.rad - yellow.asym | 0.76 | 0.23 | 3.30 | 0.03 |
| yellow.rad - yellow.asym | 0.81 | 0.23 | 3.46 | 0.02 |
| stripe.birad - black.birad | 1.85 | 0.22 | 8.58 | < 0.001 |
| yellow.birad - black.birad | 2.01 | 0.21 | 9.40 | < 0.001 |
| black.rad - black.birad | -0.14 | 0.25 | -0.55 | 1.00 |
| stripe.rad - black.birad | 1.94 | 0.23 | 8.42 | < 0.001 |
| yellow.rad - black.birad | 1.99 | 0.23 | 8.51 | < 0.001 |
| yellow.birad - stripe.birad | 0.16 | 0.22 | 0.74 | 1.00 |
| black.rad - stripe.birad | -1.99 | 0.25 | -7.94 | < 0.001 |
| stripe.rad - stripe.birad | 0.09 | 0.23 | 0.38 | 1.00 |
| yellow.rad - stripe.birad | 0.14 | 0.24 | 0.58 | 1.00 |
| black.rad - yellow.birad | -2.15 | 0.25 | -8.63 | < 0.001 |
| stripe.rad - yellow.birad | -0.07 | 0.23 | -0.30 | 1.00 |

| | | | | |
|---|---|---|---|---|
| yellow.rad - yellow.birad | -0.02 | 0.23 | -0.10 | 1.00 |
| stripe.rad - black.rad | 2.08 | 0.26 | 7.90 | < 0.001 |
| yellow.rad - black.rad | 2.13 | 0.27 | 7.99 | < 0.001 |
| yellow.rad - stripe.rad | 0.05 | 0.25 | 0.19 | 1.00 |
